# Delayed implantation induced by letrozole in mice

**DOI:** 10.1101/2021.09.15.460438

**Authors:** Fang Wang, Shijie Li, Lingshuai Meng, Ye Kuang, Zhonghua Liu, Xinghong Ma

## Abstract

Implantation timing is key for a successful pregnancy. Short delay in embryo implantation caused by targeted gene ablation produced a cascading problem in the later stages of the pregnancy. Although several delayed implantation models have been established in wild mice, almost none of them is suitable for investigating the delay on the late events of pregnancy. Here, we report a new delayed implantation model established by the intraperitoneally administration of letrozole at 5 mg/kg body weight on the day 3 of pregnancy. In these mice, initiation of implantation was induced at will by the injection of estradiol (E2). When the estradiol (3 ng) was injected on day 4 of pregnancy (i.e., without delay), the embryo implantation restarted, and the pregnancy continued normally. However, high dose of estrogen (25 ng) caused compromised implantation. We also found that only 67% of the female mice could be pregnant normally and finally gave birth when the injection of estradiol (3 ng) was on day 5 of pregnancy (i.e., one day delay). Most of the failed pregnancies had impaired decidualization, decreased plasma progesterone levels and compromised angiogenesis. Progesterone supplementation could rescue decidualization failure in the mice. Collectively, we established a new model of delayed implantation by letrozole, which can be easily used to study the effect and mechanisms of delay of embryo implantation on the progression of late pregnancy events.

## Introduction

Implantation is a process in which the embryo adheres to and invades the uterine wall, establishing physical and physiological connections with maternal endometrium for further embryonic development. Successful implantation necessitates synchronization of embryo development to the implantation competent blastocyst stage and uterine differentiation to the receptive phase[1]. Uterine receptivity for embryo implantation only lasts for a short period of time, known as the window of implantation. It is the very time that the embryo can implant into the uterus. Coordinated actions of maternal estrogen and progesterone (P4) via their cognate nuclear estrogen receptor (ER) (ERα and ERβ) and progesterone receptor (PR) (PR-A and PR-B) create the receptivity of the uterus for implantation[2]. Of the receptors, ERα (Esr1) and PR-A are important for uterine receptivity and implantation[3–5]. In mice, different uterine cell types, such as epithelial, stromal and muscle cells, respond to fluctuating estrogen and P4 levels differentially.

On day 1 of pregnancy (vaginal plug), the preovulatory secretion of estrogen stimulate epithelial cell proliferation. Reduction of ovarian hormone levels on day 2 leads to apoptosis of epithelial cells. On day 3, increased P4 secretion from the newly formed corpora lutea (CL) initiates stromal cell proliferation. On day 4, preimplantation ovarian estrogen surge superimposed with P4 further stimulates stromal cells proliferation with epithelial cell differentiation, leading to uterine receptivity for implantation. Upon embryo attachment and invasion, the stromal cells were gradually transformed to secretory decidual cells (decidualization)[1, 6].

Implantation quality determines the pregnancy outcome. A short delay of implantation (deferred implantation) caused by targeted gene deletion, such as Pla2g4a, Lpar3, Klf5 or Msx1 missing in mice, created a negative ripple effects on the late events of pregnancy leading to poor pregnancy outcome[7–10]. In human, late implantation may cause miscarriage[11]. At present, several different protocols have been used to induce delayed implantation in wide type mice and rats. Ovariectomy on the morning of day 4 of pregnancy, before the preimplantation ovarian estrogen surge, is the most commonly used experimental method to induce delayed implantation[12]. In this model, embryo implantation is postponed for a certain period, the uterus remains in a quiescent state, and blastocyst undergoes dormancy but retains its ability to initiate implantation [6, 13]. Delayed implantation can be maintained for many days by daily injection of P4, but the delayed state can be terminated by single E2 injection, and embryo implantation is initiated [14]. Delayed implantation can also be obtained by hypophysectomy in rat until the time estrogen plus progesterone or prolactin injected, or by autografts of the hypophysis[15, 16]. Non-surgical methods that block estrogen’s action or synthesis are also used to induce delayed implantation. Tamoxifen, a selective estrogen receptor modulator, is usually used to treat breast cancer. Administration of tamoxifen with progesterone resulted in the delay of implantation in rats and mice[17]. In addition, delayed implantation can be induced by antiestrogenic compounds MER-25 or 10275-S, or by an estrogen receptor antagonist (ICI 182,780) in rats[18].

Aromatase is a cytochrome P450 enzyme responsible for the last step in estrogen biosynthesis, catalyzing the aromatization of androstenedione and testosterone into estrone and estradiol, respectively **[19]**. Aromatase inhibitors can prevent estrogen production by inhibiting aromatase activity **[20]**. Although several aromatase inhibitors have been reported to have the ability to inhibit E2 and disturb embryo implantation in rats by mini-osmotic pump **[21–23]**, it is unable to exactly control the function time of E2. Therefore, a better delayed implantation model is needed. Letrozole is a third-generation aromatase inhibitor and shows greater selectivity and efficacy over old agents **[24, 25]**. It is widely used in treatment of human disease that caused by abnormal estrogen level, such as breast cancer, ovulation induction and endometriosis **[19, 26, 27]**. We are wondering whether letrozole can be used to induce delayed implantation to explore the effect of the delay on the development of pregnancy in late stage.

In this study, we established a new model of delayed implantation by using letrozole to inhibit pre-implantation E2 surge and postponed the time of E2 exposure by controlling exogenous E2 injection time. We found that decidualization was failed and pregnancy was lost in the 1-day delayed implantation mice. Decrease of P4 level was regarded as one of the reasons for this failure, and exogenous supplement of P4 could rescue this decidualization process.

## Materials and methods

### Animals

All experiments were approved by Animal Care and Use Committee of the Northeast Agricultural University. Mice were housed under a 12-hour light-dark cycle condition and with water and food available <u>ad libitum</u>. Female ICR mice (8~10 weeks old) were mated with fertile males, and the day of vaginal plug was defined as day 1 of pregnancy.

### Model of delayed implantation induced by letrozole

In order to explore the optimal dose of delayed implantation, letrozole (Tocris) was dissolved in DMSO in a dose of 0.1, 0.5, 0.8, 1, 2, and 5 mg/kg body weight, respectively. Mice were injected intraperitoneally with different dose of letrozole at 1200 h on day 3 of pregnancy, and control mice were injected with 50 μl DMSO alone. Implantation sites were checked on day 5 by intravenous blue dye injection. In order to exclude the situation of failed fertilization, uteri from these mice with no blue dye reaction were collected and flushed with 0.9% saline water. Only the mice with embryos were counted as delayed implantation.

In the 0-day delayed model, delayed implantation was induced by injection with letrozole at a dose of 5 mg/kg body weight on the day 3 of pregnancy, and 3 ng E2 or 25 ng E2 was injected on the morning of day 4 to activate embryo implantation.

In the 1-day delay model, delayed implantation was induced by injection with letrozole at a dose of 5 mg/kg body weight on the day 3 of pregnancy, and 3 ng E2 was injected subcutaneously on the morning of day 5 to activate embryo implantation.

### Model of ovariectomy which induced delayed implantation

We established the traditional delayed implantation mice based on previous reports [14]. In brief, the ovaries of mice were removed on the morning of day 4, and then exogenous P4 was injected to maintain the delayed state and 25 ng E2 was injected subcutaneously to restart embryo implantation.

### Hematoxylin and eosin (H&E) staining

Paraffin embedded tissues were sectioned at 5 μm thick and dried at 37°C overnight. Then the sections were deparaffinized by xylene twice for 5 min each. For rehydration, the sections were washed in 100%ethanol twice for 5 min each, 95% ethanol for 2 min, 70% ethanol for 2 min and rinsed in distilled water. After staining with hematoxylin stain for 5 min, the sections were differentiated in 1% acid alcohol for 30 sec and blued in 0.2% ammonia water for 1 min. Then the sections were rinsed in 95% ethanol for 10 sec and counterstained in the eosin stain for 1 min. Finally, the sections were dehydrated with ascending alcohol solutions, cleared in xylene and mounted with Permount.

### Immunohistochemistry

Tissues were fixed in 10% formalin solution and embedded in paraffin. Sections at 5 μm thick were deparaffinized by xylene and rehydrated in a graded alcohol series. After blocking by 10% horse serum for 1 h at RT, sections were incubated by PECAM antibodies (R&D Systems, AF3628) at 4°C overnight. On the second day, sections were washed by PBS and incubated by HRP conjugated goat anti-rabbit IgG secondary antibody (Vector laboratory, BA-1000) for 1 h at RT followed by incubation with horseradish peroxidase conjugated streptavidin (Vector laboratory, SA-5004). Positive signals were detected by DAB (Sigma-Aldrich, D8001).

### In situ hybridization

For probe construction, total RNA was extracted from mouse uterus and reverse transcribed into cDNA by reverse transcriptase kit (Promega). cDNA was amplified by gene specific primers. PCR products were cloned into pGEM-T vectors. cRNA probes were generated by Digoxin labeling kit (Roche) according to the manufacturer’s instructions. For hybridization, frozen sections at 10 μm thick were fixed in 4% paraformaldehyde for 1 h at room temperature (RT). The sections were treated with Triton X-100 at RT for 20 min. After hybridized in the solution of 50% formamide hybridization buffer containing sense or antisense cRNA probes at 55°C overnight, slides were blocked with 1% blocking reagent (Roche) for 1 h at RT. Then the sections were incubated with anti-Digoxin antibody conjugated with alkaline phosphatase (Roche) at 4°C overnight. The signal was detected after incubation in NBT/BCIP solution (Roche) for an optimal time. Primer sequences used for producing cRNA probes were listed as follows: *COX-2*, forward, 5’-CCCCCACAGTCAAAGACACT-3’, reverse, 5’-GAGTCCATGTTCCAGGAGGA-3’; *Hoxa10*, forward, 5’-CCCACAACAATGTCATGCTC-3’, reverse, 5’-TGTTCTGCGCAAAAGAACAC-3’; *Ihh*, forward, 5’-CTCACTGGCCATCTCTGTCA-3’, reverse, 5’-CTCGATGACCTGGAAAGCTC-3’.

### Measurement of Serum P4 Level

In order to check the P4 level during decidualization in the 0-day delay mice and 1-day delay mice, we collected blood samples from the mice of 0-day delay and 1-day delay model at 36 h later since E2 injection. The blood samples were stored at room temperature for 2 h. Serum was separated by centrifugation and then stored at −70°C for further analyses. The P4 levels in the serum were measured using commercial Kit (Roche).

### Statistical analysis

Data are expressed as mean ± SEM. The significance of differences was analyzed by Unpaired Student’s *t*-test using the GraphPad Prism 5 (GraphPad Software, CA). *P* < 0.05 was considered significantly different.

## Results

### Establishment of a new model of delayed implantation by letrozole

In mice, embryo implantation occurs under the function of a small amount of ovarian estrogen on the morning of day 4 [12]. To establish the delayed implantation model, we tried to inhibit E2 production by single letrozole injection. First, we explored the optimal dose of letrozole in inducing delayed implantation. We checked the effect of letrozole at a dose of 0.1, 0.5, 0.8, 1, 2, and 5 mg/kg body weight, respectively. As shown in Fig. 1a, low dose of letrozole could slightly inhibit embryo implantation, since weak blue bands in uteri could be observed. Medium dose of letrozole such as 1 mg and 2 mg could partially delayed implantation. Some mice showed weak blue dye reaction, but some mice showed no blue dye reaction. We found that letrozole at a dose of 5 mg/kg body weight could completely inhibit embryo implantation in all the examined mice (n=5). Blastocyst were recovered from the delayed uteri (Fig. 1b). Therefore, we chose the dose of 5 mg/kg body weight as the optimal dose to induce delayed implantation in our study. Then we tried to activate embryo implantation by E2 injection following the letrozole injection. We found that both 3 ng and 25 ng E2 could normally restart embryo implantation when injected on day 4 in delayed implantation mice induced by letrozole (Fig. 1c and d). The mice had normal number of implantation sites compared to the control mice that injected with DMSO (Fig. 1e).

**Fig. 1.**
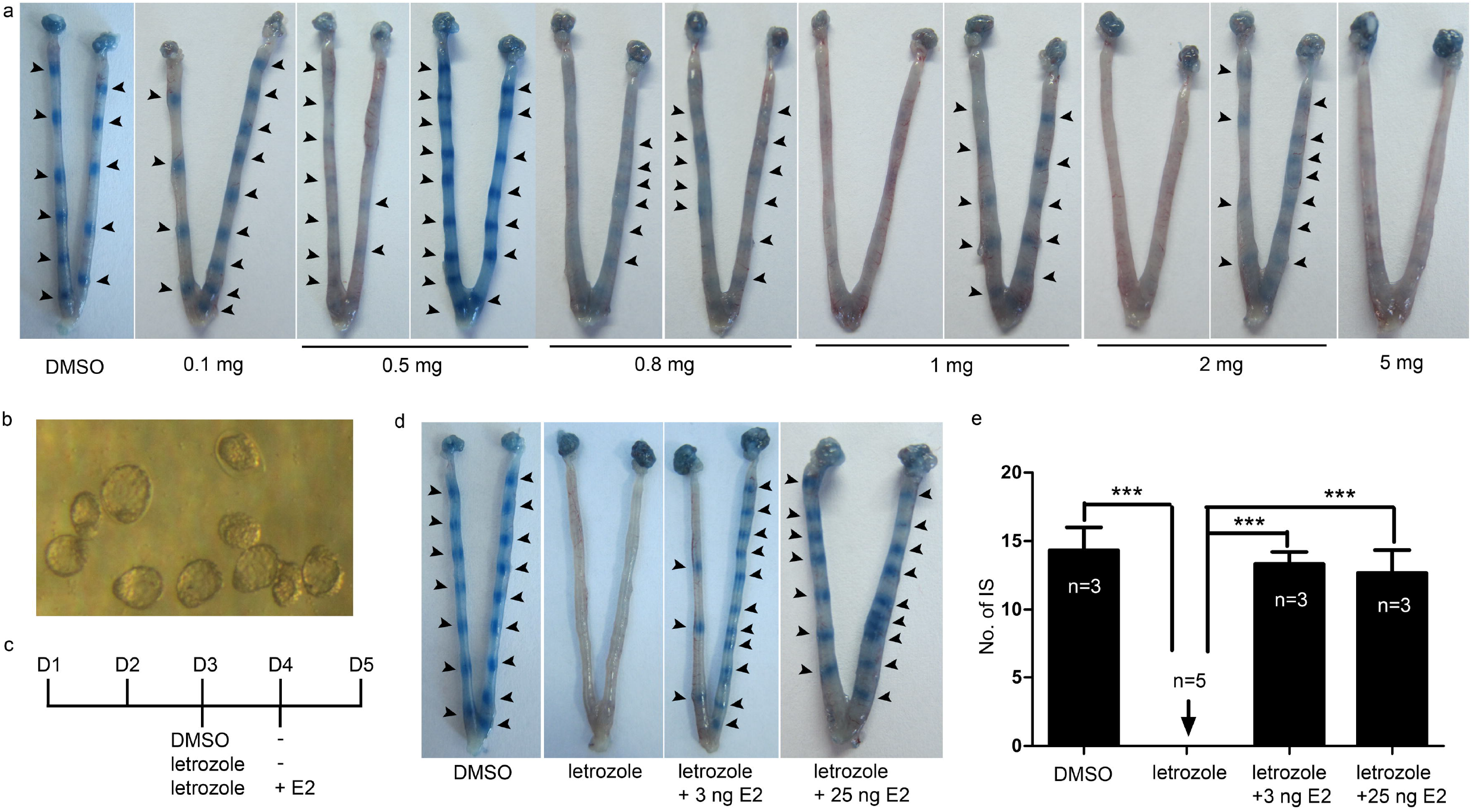
Model of letrozole induced delayed implantation. (a) Delayed implantation. Representative mice of delayed implantation induced by letrozole at a dose of 0.1, 0.5, 0.8, 1, 2, and 5 mg/kg body weight, mice injected with DMSO as control. (b) Blastocysts recovered from the uteri of delayed implantation mice. (c) Experimental scheme. To delayed implantation, mice received letrozole treatment on day 3. To restart implantation, letrozole treated mice received 3 ng or 25 ng E2 on the morning of day 4. Mice only received DMSO on day 3 were regarded as control. (d) Activated implantation. Representative of activated implantation by 3 ng or 25 ng E2 in mice of letrozole induced delayed implantation. (e) Number of implantation sites (IS) in (d). Numbers within columns indicate the total number (n) of mice examined. Results are shown as mean ± SEM (Tukey’s test, \*\*\**P* < 0.0001). Arrowheads indicate the implantation site

### 25 ng estrogen cause compromised implantation in the 1-day delay model, P4 supplementation could rescue the failure

In our study, we tried to explore the effect of delayed implantation for 24 h on the pregnancy. As shown in Fig. 1a, single injection with 5 mg/kg body weight letrozole was sufficient to delay implantation for one day. Next, we explored the optimal dose of E2 in activating embryo implantation in the 1-day delay model. To restart implantation in letrozole treated mice, 3 ng or 25 ng E2 was injected on day 5, implantation sites were checked on the morning of day 6 (Fig 2a). As shown in Fig. 2b and 2d, all the mice that received 3 ng E2 restarted implantation normally (100%, n=7). By contrast, mice that received 25 ng showed low implantation rate (43%, n=7). Only 2 out of 7 mice that received 25 ng E2 showed normal implantation, one mouse had weak blue dye reaction, and 4 out of 7 mice showed no implantation sites. Blastocysts were recovered from uteri with no or weak blue dye reaction (Fig. 2c). It is widely accepted that E2 and P4 are the two major regulators in reproduction that usually show antagonistic effect [28]. We explored to rescue this compromised implantation by exogenous supplement with P4. We observed that all the mice (n=6) showed normal blue dye reaction after a single injection of 1 mg P4 (Fig. 2b).

**Fig. 2.**
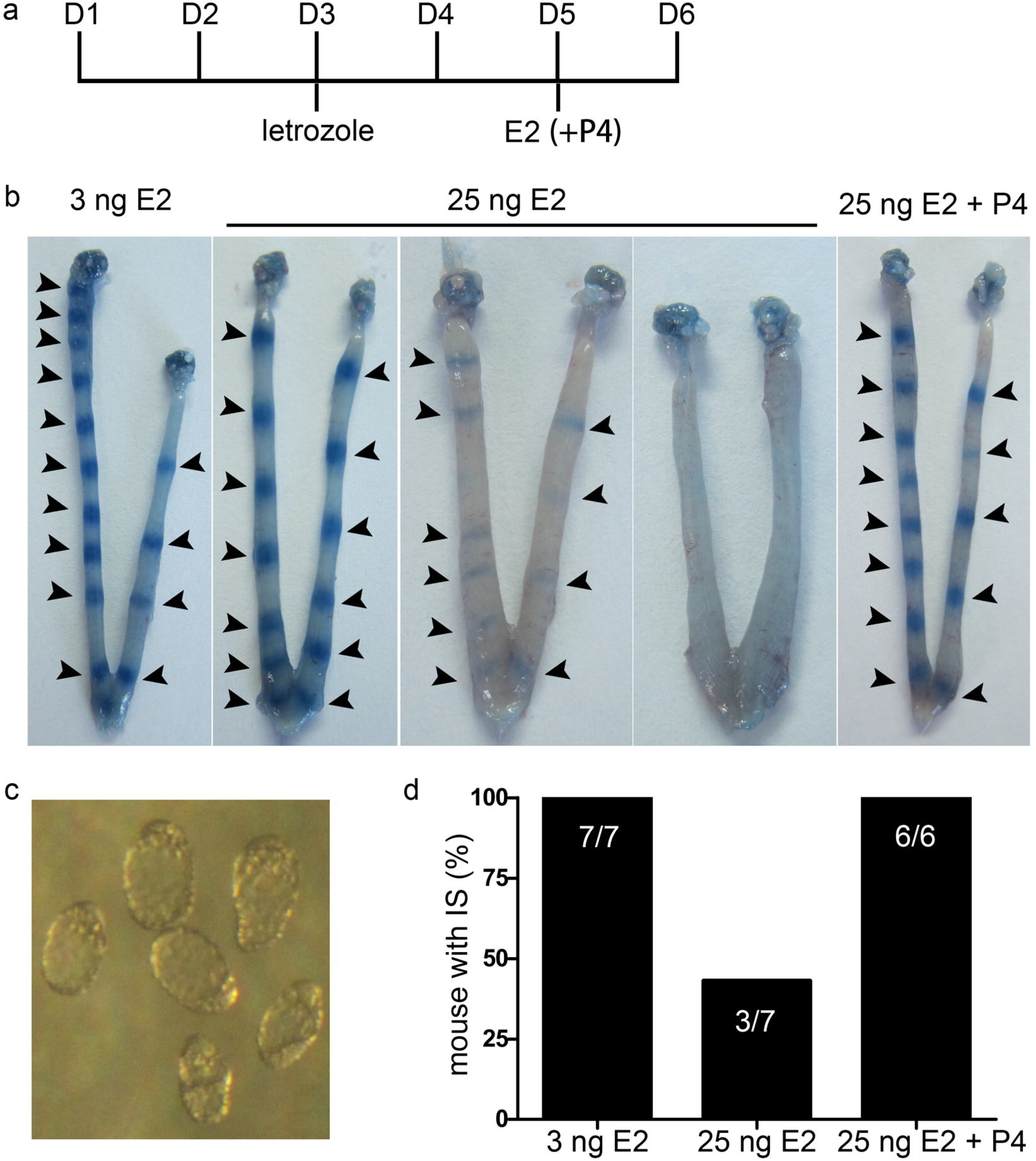
25 ng E2 causes compromised implantation in 1-day delay model. (a) Experimental scheme. Letrozole treated mice were injected with 3 ng, 25 ng E2 or 25 ng E2 plus 1 mg P4 on the morning of day 5, uteri were collected on day 6. (b) Representative mice that received 3 ng E2, 25 ng E2, and 25 ng E2 combined with P4 in 1-day delay model. Implantation sites were examined on day 6 by blue dye reaction. Arrowheads indicate the implantation sites. (c) Uteri of weak or no blue dye in (b) were flushed and blastocysts were shown. (d) Percentage of mice with implantation sites (IS)/total mice examined. Numbers within columns indicate the total number (n) of mice examined

### 25 ng E2 cause abnormal expression of implantation related genes in the 1-day delay model

Next, we explored the reasons for the 25 ng E2 caused implantation failure. First, we checked the expression of *Ihh* and *Hoxa10* in preimplantation uteri. The *Ihh* gene was expressed in the uterine epithelia, and showed the same expression pattern in all mice examined (Fig. 3). Whereas the mRNA of *Hoxa10* in uterine stromal cells decreased to a very low level in the mice injected with 25 ng E2. Next, we checked the expression pattern of *COX-2* in embryo implantation sites. Similarly, the expression of *COX-2* mRNA decreased hardly around the implantation sites in the 1-day delay mice treated with 25 ng E2. However, P4 supplementation could rescue the expression of *Hoxa10* and *COX-2* in the 1-day delay mice treated with 25 ng E2. These results suggest that 3 ng is the optimal dose of E2 to initiate implantation in our 1-day delay model, which was chosen for further experiments.

**Fig. 3.**
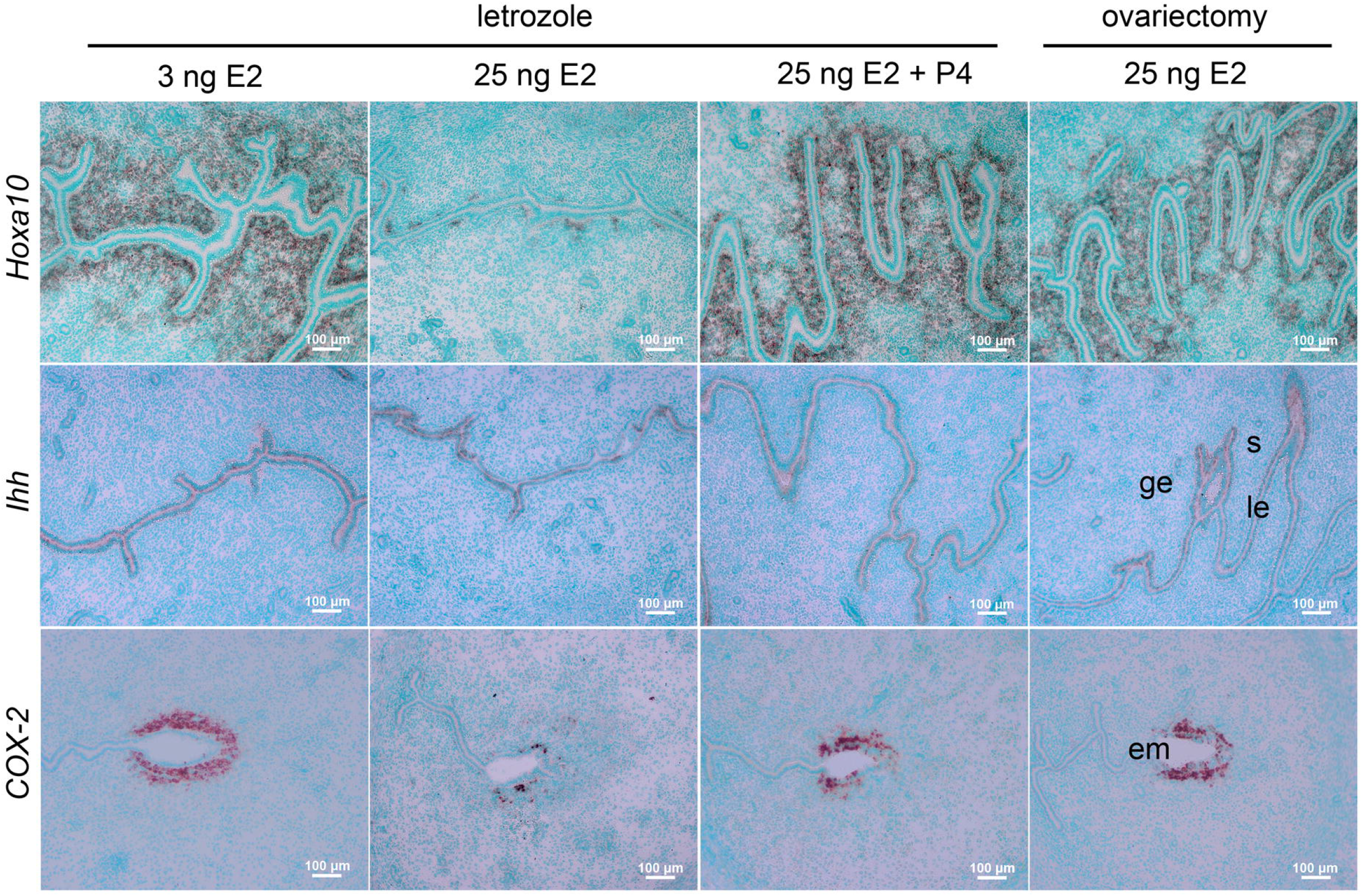
25 ng E2 causes abnormal expression of implantation related genes. In situ hybridization of *Hoxa10, Ihh* and *COX-2* in the 1-day delay mice activated by 3 ng E2, 25 ng E2, 25 ng E2 combined with P4, and traditional ovariectomized mice activated by 25 ng E2. Le, luminal epithelium; ge, glandular epithelium; s, stroma; em, embryo. Bars represent 100 μm

### Comparison between traditional ovariectomized model and letrozole induced delayed implantation model

In traditional delayed implantation mice, 25 ng E2 can start implantation. In our 1-day delay model, 3 ng E2 could start implantation, but 25 ng E2 caused implantation failure. We examined the expression of hormone regulated genes *Ihh* and *Hoxa10* and angiogenesis associated gene *COX-2*, which are important genes for successful implantation [29–31], in these mice. The in-situ hybridization results showed that the expression patterns of these genes in the 1-day delay mice treated with 3 ng E2 was similar to that in the traditional delayed implantation mice treated with 25 ng E2 (Fig. 3). Under the same E2 level (25 ng E2), the different gene expression patterns between the letrozole induced delayed implantation model and the traditional one suggests that the two models are different in E2 sensibility.

### Pregnancy outcome in 1-day delay model

Many previous researches had shown that deferred implantation led to pregnancy loss. We explored the pregnancy of the mice in our letrozole induced delayed implantation model. As shown in Table 1, both control (DMSO) mice and 0-day delay mice had normal pregnancy rate. However, mice in 1-day delay model showed comprised pregnancy outcome. Only 67% mice could give birth to pups, whereas the number of pups delivered was normal, approximately 12 pups per mouse. The results show that the implantation postponed for one day by E2 cause compromised pregnancy outcome.

**Table 1.** Pregnancy outcome in mice of delayed implantation induced by letrozole

| | No. Dams | No. Delivered dams | Pregnancy rate (%) | No. Pups per dam (mean ± SEM) |
| --- | --- | --- | --- | --- |
| Control | 7 | 7 | 100 | 13.6 ± 1.6 |
| 0-day delay | 10 | 9 | 90 | 12.1 ± 3.7 |
| 1-day delay | 12 | 8 | 67 | 12.1 ± 2.3 |

Pregnancy rate was calculated by No. Delivered dams/total dams. There was no difference in number of pups between 0-day delay and 1-day delay mice.

### Compromised angiogenesis in 1-day delay model

Angiogenesis is an important event that support the growth of the implanted embryos during the decidualization process [32]. Here, we examined the angiogenesis in our delayed model. We checked the vascular permeability in mice from 0-day delay and 1-day delay model at 36 h and 48 h later since E2 injection by blue dye reaction (Fig 4a). As shown in Fig. 4b and c, mice of 0-day delay model showed normal blue dye reaction at the both time points. However, mice of 1-day delay model showed compromised vascular permeability. At 36 h after E2 injection, we observed that one mouse showed normal implantation, 2 out of 5 mice showed weak blue dye reaction, and the other two mice had no blue dye reaction completely. The blastocysts were recovered from the mice that had no blue dye reaction. At 48 h after E2 injection, 3 out of 7 mice showed normal decidualization, four mice showed compromised decidualization: one mouse had normal number of implantation sites with no blue dye reaction; two mice showed “white band” in uteri; one mouse had no implantation sites, while corpus luteum in the ovary suggested the pregnancy occurred. PECAM is a cellular adhesion and signaling receptor, which plays an important role in regulation of the vascular permeability barrier during angiogenesis [33, 34]. To further explore the angiogenesis in 1-day delay mice, we checked the expression of PECAM at 48 h later since E2 injection. As shown in Fig 4d, mice from pregnant day 6, 0-day delay model and 1-day delay model with normal blue dye reaction showed similar PECAM expression pattern. However, mice with abnormal blue dye reaction of 1-day delay model showed markedly reduced PECAM staining in the stromal region of the uteri. As increased vascular permeability is necessary for formation of decidua structure, we next checked the structure of decidua in 0-day delay and 1-day delay model at 48 h later after E2 injection by H&E staining. As shown in Fig 4e, there are three types of decidualization at the implantation sites in the 1-day delay model: some mice showed normal decidualization compared to that in 0-day delay model, some mice formed the “tumor” like structure, and some mice showed implantation sites resorbed completely. These results suggested that the decidualization could initiate normally at early stage in the 1-day delay model, while the subsequent decidual response in some mice was gradually attenuated and some of implantation sites were resorbed due to compromised vascular permeability. Collectively, these results indicated that compromised angiogenesis occurred during decidualization process in mice that E2 was postponed for 24h.

**Fig. 4.**
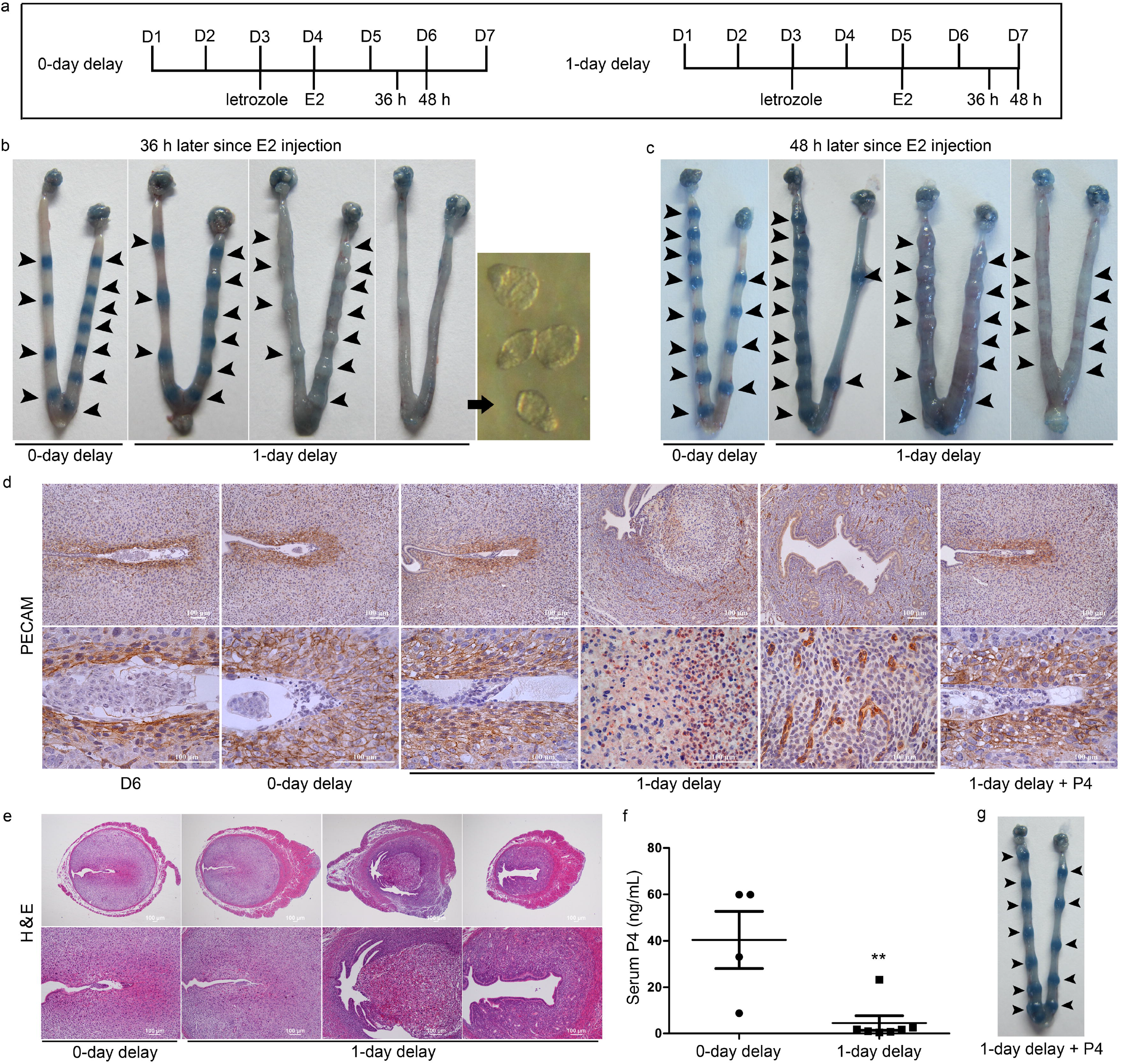
Angiogenesis in 1-day delay mice. (a) Experimental scheme of 0-day delay and 1-day delay model. (b) Representative mice at 36 h since E2 injection in the 0-day delay and 1-day delay model. Arrow indicates the blastocyst recovered from the uteri that show weak or no blue dye reaction. Arrowheads indicate the implantation sites. (c) Representative mice at 48 h later since E2 injection in the 0-day delay and 1-day delay model. (d) Immunohistochemistry of PECAM. PECAM immunostaining was performed in mice of 0-day delay and 1-day delay treated with or without P4 mice at 48 h since E2 injection. Normal pregnant mice on day 6 were regarded as control. (e) H&E staining of implantation sites and resorption sites in (c). (f) Serum P4 levels at 36 h later since E2 injection in the 0-day delay mice and 1-day delay mice. Statistical analysis was done by unpaired *t*-test, \*\**P* < 0.01. (g) Representative uteri on day 7 from the 1-day delay mice that supplemented with 1 mg P4. Mice from 1-day delay model received single P4 supplementation at the time of E2 injection on day 5 and then uteri were collected on the morning of day 7 after intravenous injection of trypan blue solution. Arrowheads indicate the implantation sites.

### Delayed implantation induced by letrozole cause rapid decrease of serum P4, supplement with P4 could rescue angiogenesis

It is interesting to know the mechanism of angiogenesis failure in 1-day delay mice. Several reports have shown that P4 plays a role in endometrial angiogenesis [35–37]. Otherwise, it is known that P4 levels are rapid decreased on day 6 of pseudopregnancy in mice due to luteolysis [38]. Therefore, we analyzed the level of serum P4 in 0-day delay mice and 1-day delay mice. As shown in Fig. 4f, serum P4 level decreased rapidly in 1-day delay mice compared with 0-day delay mice at 36 h after E2 injection. To explore whether P4 supplementation could rescue the failed angiogenesis in 1-day delay model, 1 mg P4 was injected into the letrozole induced delayed-implantation mice at the time of E2 administration on day 5. The result showed that a single injection with 1 mg P4 could rescue decidualization (Fig. 4g). All the mice (n=6) received single P4 injection showed normal blue dye reaction when checked on the morning of day 7. Otherwise, the expression pattern of PECAM was also restored into normal state in pregnant mice (Fig 4d). These results suggested that decreased P4 level was one of the reasons of compromised angiogenesis in 1-day delay mice.

## Discussion

Implantation timing is key for successful pregnancy. In order to investigate the effect of delayed implantation on pregnancy progression and outcome and the mechanism, we established a new delayed implantation model, in which single injection of letrozole at a dose of 5 mg/kg body weight could completely inhibit implantation and single estrogen injection could restart implantation. At present, ovariectomy is the most commonly used method to establish delayed implantation. This method is time-consuming and places much stress on experimental animals. Especially, after removing the ovaries, it becomes complex and difficult to study the development of late pregnancy because the progression of pregnancy depends largely on ovarian estrogen and progesterone [37]. Our method is easy to operate and can effectively induce the delay of implantation. In our model, delay of implantation and reactivation of implantation can be achieved by one injection of letrozole and estrogen, respectively. In this model, the use of letrozole does not affect the subsequent development of pregnancy evidenced by normal implantation, decidualization and pregnancy outcome in females of 0-day delay model in which the exogenous estrogen is timely supplemented on the fourth day of pregnancy (i.e., without implantation delay). However, the delay of implantation for one day by postponed E2 injection for one day caused decidualization failure and compromised pregnancy. Similar phenomenon is observed in targeted gene deletion mice, in which the deferred implantation had negative ripple effect on post-implantation events and lead to poor pregnancy outcome [5, 7]. These results indicated that timely implantation is necessary for successful pregnancy. Additionally, we found that serum P4 level in 1-day delay model decreased compared to that in 0-day delay mice, and supplement with P4 could rescue the decidualization process. Therefore, we speculated that decrease of P4 level was one of the reasons for decidualization failure in the 1-day delayed mice.

In our 1-day delay model, letrozole administration and delayed implantation caused rapid decrease of P4. Similar phenomenon was observed in letrozole induced polycystic ovaries in rats, whose P4 level was decreased after daily administration of letrozole [39]. It is reported that E2 has the luteotropic function [40, 41], and the inhibition of E2 synthesis with an aromatase inhibitor resulted in the reduction of serum P4 level in pseudopregnant rats [42]. However, in our 0-day delay model, the exogenous E2 was supplied at the time when endogenous ovarian E2 surge occurred, and P4 level was comparable to the control mice. These results suggested that timely E2 surge on the morning of day 4 or the timely on-time implantation itself might be necessary for luteal P4 secretion. Alternatively, it is presumed that E2 might increase the availability of cholesterol substrate for P4 synthesis by regulating lipoprotein receptor content or stimulating cholesterol uptake, intra-cellular transport and synthesis [43]. To further understand the mechanism of P4 decrease in our model, future experiments might be needed to investigate the changes of cholesterol levels after letrozole treatment.

The progesterone P4 is critical for developmental events during the pregnancy, and P4 levels vary throughout the pregnancy [44]. The P4 level required for pregnancy maintenance is higher than early developmental events such as implantation and decidualization [45]. In our model, single P4 injection could only restore the decidualization until day 7, and some mice showed compromised decidualization again on day 8. To rescue late developmental events in our 1-day delay model, larger amount of P4 was required. We found that post-implantation events were restored in mice with daily P4 supplementation when checked on day 13. These results suggested that insufficient P4 level is the major cause of decidualization failure and poor pregnancy outcome of our 1-day delayed implantation model.

E2 plays an important role in regulating the implantation window. Of the women that received embryo transfer, the pregnant women have a lower E2 level compared to the non-pregnant women [46]. In the traditional delayed implantation model, low level of E2 (3 ng) made the implantation window remain open for an extended period, while high level of E2 (25 ng) rapidly close the window [47]. Whereas, both the E2 levels could activate the dormant blastocysts. In our 1-day delay model, in contrary to the traditional model, 3 ng E2 could activate dormant blastocyst to implant, but 25 ng E2 compromised the initiation of implantation. Meanwhile, the genes regulated by P4, such as *Hoxa10* and *Ihh*, were downregulated in the stromal cells in our 1-day delay model. The distinct phenomenon observed in the two models could be due to the different levels of P4. The P4 level was high in the traditional ovariectomized mice because large amount of exogenous P4 was supplemented, while P4 level in the letrozole treated mice was at the physiological level. Therefore, the E2/P4 ratio is higher in our letrozole induced delayed mice than traditional ovariectomized mice.

Delayed implantation has evolved naturally across more than 100 species of mammals for dealing with an unpredictable environment and food shortage[48]. It seems that this kind of evolutionary selected delay of implantation has no adverse effects on the development of late pregnancy[49]. The implantation delay resulted from the suckling in lactating mice and rats also seems to have no bad effect on subsequent pregnancy[50, 51]. However, deferred implantation caused by targeted gene deletion had multiple negative effects on pregnancy, including fetal growth retardation, embryo crowding,embryonic death, and multiple embryos sharing one placenta[7, 8, 10, 52]. For the delayed embryo implantation induced by the traditional surgical methods, autophagy occurred in the dormant blastocysts during the delay[53]. Interestingly, delayed implantation caused by asynchronous transplantation does not affect decidualization, but has adverse effects on pregnancy[7, 54]. In our new model, however, delayed implantation affects not only the decidualization but also the development of late pregnancy. Therefore, establishment of a variety of different delayed implantation models is conducive to study the impact of early pregnancy events on the progress of late pregnancy. Our new delayed implantation model contributes to the understanding of the effect of early events on the development of late pregnancy.

## Declarations

### Availability of data and material

The data and material that support the findings of this study are available from the corresponding author upon request.

### Authors’ contributions

X.M. and Z.L. conceived and designed research; F.W., L.M. and S.L. performed experiments; X.M., F.W. and Y.K. analyzed the data; X.M. and F.W. wrote the manuscript.

